# Microtubules promote intercellular contractile force transmission during tissue folding

**DOI:** 10.1101/540344

**Authors:** Clint S. Ko, Vardges Tserunyan, Adam C. Martin

## Abstract

During development, forces transmitted between cells are critical for sculpting epithelial tissues. Actomyosin contractility in the middle of the cell apex (medioapical) can change cell shape (e.g., apical constriction), but can also result in force transmission between cells via attachments to adherens junctions. How actomyosin networks maintain attachments to adherens junctions under tension is poorly understood. Here, we discovered that microtubules promote actomyosin intercellular attachments in epithelia during *Drosophila* mesoderm invagination. First, we used live imaging to show a novel arrangement of the microtubule cytoskeleton during apical constriction: medioapical Patronin (CAMSAP) foci formed by actomyosin contraction organized an apical non-centrosomal microtubule network. Microtubules were required for mesoderm invagination but were not necessary for initiating apical contractility or adherens junction assembly. Instead, microtubules promoted connections between medioapical actomyosin and adherens junctions. These results delineate a role for coordination between actin and microtubule cytoskeletal systems in intercellular force transmission during tissue morphogenesis.

## Introduction

Apical constriction is a ubiquitous cell shape change that results in dramatic rearrangements of tissue architecture, such as tissue folding (Sawyer et al., 2010; Heisenberg and Bellaïche, 2014; Martin and Goldstein, 2014). The cellular force necessary to constrict a cell apex is generated by actomyosin contraction, which is regulated by RhoA signaling (Jaffe and Hall, 2005; Kasza and Zallen, 2011; Lecuit et al., 2011). During apical constriction, the apical cortex is often polarized; myosin-II (myosin) is activated near the middle of the apical cortex (medioapical), which contracts an actin filament (F-actin) network that spans the apical surface (Sawyer et al., 2009; Blanchard et al., 2010; David et al., 2010; Mason et al., 2013; Booth et al., 2014; Sánchez-Corrales et al., 2018). In order for these changes in cell geometry to cause tissue morphogenesis, cellular forces must be transmitted and integrated across the tissue (Fernandez-Gonzalez et al., 2009; Lecuit and Yap, 2015). This is mediated by connecting contractile actomyosin meshworks to E-cadherin-based adherens junctions (Martin et al., 2010, Sawyer et al., 2011). Molecular components that mediate this linkage have been identified and are important for morphogenesis (Sawyer et al., 2009; Desai et al., 2013). In addition, this attachment has been shown to be dynamic and actin turnover is required to promote attachment by repairing lost connections (Roh-Johnson et al., 2012; Jodoin et al., 2015). However, whether other mechanisms maintain actomyosin network connections to junctions, in the face of tension, remains unknown.

During gastrulation in the early *Drosophila* embryo, apical constriction leads to mesoderm and endoderm cell invagination (Leptin and Grunewald, 1990; Sweeton et al., 1991) (Fig. 1 A). Mesoderm cells express transcription factors, Twist and Snail, that promote apical RhoA activation, which induces actomyosin contractility (Barrett and Settleman, 1997; Häcker and Perrimon, 1998; Dawes-Hoang et al., 2005; Fox and Peifer, 2007; Kölsch et al., 2007; Izquierdo et al., 2018). Contractile force is transmitted across the folding tissue through adherens junctions, resulting in epithelial tension predominantly along the anterior-posterior axis (Martin et al., 2010; Chanet et al., 2017). Apical constriction in multiple invagination processes depends on polarized RhoA signaling, with active RhoA and its downstream effector Rho-associated coiled-coil kinase (ROCK), which activates myosin (Amano et al., 1996; Mizuno et al., 1999), being enriched in the middle of the apical surface (Mason et al., 2013; Booth et al., 2014; Coravos et al., 2016; Chung et al., 2017). It is poorly understood how intercellular actomyosin connections are promoted when the medioapical pool of active RhoA is present at a distance from cell junctions.

**Figure 1.**
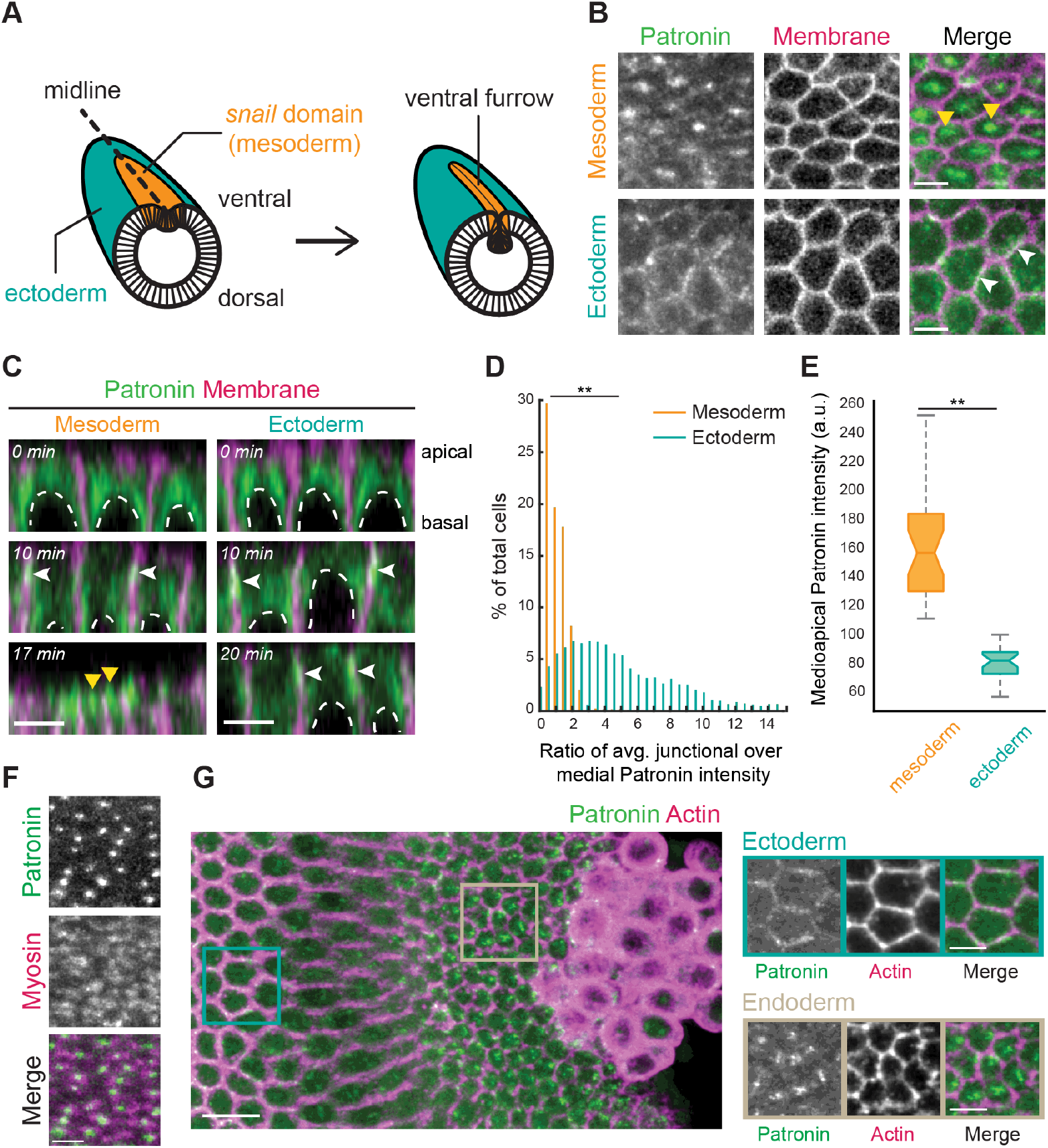
Patronin::GFP localizes medioapically in apically constricting cells. (A) Diagram of an embryo undergoing mesoderm invagination. Ventral, mesoderm cells (*snail* expressing domain highlighted in red) apically constrict and internalize, forming a ventral furrow along the midline (dashed line). (B) Patronin::GFP is present in a medioapical focus specifically in the mesoderm (top row, yellow arrowhead). Patronin::GFP is enriched at junctions in the ectoderm (bottom row, white arrowhead). Images are maximum intensity projections from a live embryo expressing Patronin::GFP (apical surface) and Gap43::mCH (plasma membranes, sub-apical slice). (C) Patronin::GFP localization changes from junctional (white arrowheads) to medioapical (yellow arrowheads) in the mesoderm. Images are apical-basal cross-sections from a live embryo expressing Patronin::GFP and Gap43::mCH. Top row: mid-cellularization; middle row: late-cellularization/early gastrulation; bottom row: during folding. Nuclei are highlighted by dashed white lines. (D) Quantification of medioapical Patronin::GFP enrichment. Individual cells were segmented and the junctional and medioapical Patronin::GFP intensity was calculated and the distribution of the ratio (junctional/medioapical) was plotted as a percentage of cells within each bin (n = 6 embryos, 559 cells; **, P <.0001, Kolmogorov-Smirnov test). (E) Apical Patronin::GFP foci are more intense in the mesoderm than the ectoderm. The maximum apical Patronin::GFP intensity was determined in a region encompassing the medioapical cortex in both the mesoderm (left) and ectoderm (right) (n = 6 embryos, 10 measurements per embryo; **, P < .0001, unpaired t test). Notch is the median, bottom and top edges of the box are the 25^th^ and 75^th^ percentiles, whiskers extend to the most extreme data points. (F) Medioapical Patronin::GFP co-localizes with apical myosin patches. Images are apical surface Z-projections from a live embryo expressing Patronin::GFP and Myo::mCH (sqh::mCH). (G) Patronin::GFP localizes medioapically in apically constricting endoderm cells. Images are maximum intensity projections of a fixed embryo expressing Patronin::GFP (apical surface). The embryo was immunostained with Phalloidin conjugated with AF647 to visualize cell outlines (sub-apical section). Scale bars, 10 μm (**G, left**) and 5 μm (**B-F; G, right**).

While the regulation and organization of actomyosin during apical constriction has been well-studied, the organization of the microtubule cytoskeleton and its importance is less well understood. In epithelia, there is evidence for different microtubule functions, such as regulating assembly or position of the adherens junctions (Harris and Peifer, 2005; Stehbens et al., 2006; Meng et al., 2008; Le Droguen et al., 2015), recruiting apical myosin (Booth et al., 2014), regulating RhoA (Rogers et al., 2004; Nagae et al., 2013), and providing a counter-force resisting actomyosin contraction (Takeda et al., 2018; Singh et al., 2018). Importantly, studies have shown that microtubules are necessary for cell shape changes, like apical constriction (Lee and Harland, 2007; Lee et al., 2007; Booth et al., 2014). However, it is unknown whether microtubules also play additional roles in promoting tissue shape changes, which requires imaging microtubules with high spatial and temporal resolution at both cell- and tissue-scales.

Here, we investigated microtubule cytoskeleton organization and its mechanistic contribution to mesoderm invagination. We showed that a microtubule minus-end-binding protein, Patronin, co-localized with myosin and that actomyosin contractility organized microtubules in the medioapical cortex of apically constricting cells. We found that microtubules were required for mesoderm invagination, but not apical cortex contractility. Instead, proper microtubule organization promoted actomyosin attachment to adherens junctions. These results suggest that crosstalk between actomyosin and microtubules is critical for apical actomyosin attachments to adherens junctions and, thus, intercellular force transmission.

## Results

### Medioapical Patronin focus co-localizes with myosin in apically constricting cells

To determine if the microtubule cytoskeleton plays a role in the apical constriction of mesoderm cells of the early *Drosophila* embryo, we first sought to establish the organization of the microtubule cytoskeleton. In most polarized epithelial cells, linear arrays of microtubules align with the apical-basal axis such that the minus ends are uniformly localized across the apical surface (Bacallao et al., 1989; Khanal et al., 2016; Toya et al., 2016). These minus ends are often capped and stabilized by a family of calmodulin-regulated spectrin-associated proteins (CAMSAPs or Patronin in *Drosophila*), which specifically bind to microtubule minus ends (Goodwin and Vale, 2010; Khanal et al., 2016; Noordstra et al., 2016; Toya et al., 2016). We imaged live and fixed *Drosophila* embryos expressing Patronin::GFP driven by the ubiquitin promoter (Ubi-p63E-Patronin::GFP, hereinafter referred to as Patronin::GFP). During gastrulation, there was a striking difference in Patronin::GFP localization between mesoderm (apically constricting) and ectoderm cells (not constricting) (Fig. 1 B). Specifically, Patronin::GFP was polarized to a central focus in the medioapical cell cortex of mesoderm cells, but was enriched at cell junctions in the ectoderm (Fig. 1, B and C; Video 1).

To demonstrate the difference in Patronin localization between these cell types, we measured the ratio of average junctional Patronin::GFP intensity to the average medial Patronin::GFP intensity. During mesoderm invagination, there was a significant difference in the ratio of junctional to medial Patronin::GFP intensity between mesoderm and ectoderm cells, with medial enrichment of Patronin::GFP intensity being highest in the mesoderm (mean junctional to medial ratio of 0.75 ± .53 compared to 4.75 ± 3.47) (Fig. 1 D). In addition, the maximum medioapical Patronin::GFP intensity was higher in the mesoderm than the ectoderm, consistent with Patronin forming a large apical focus in these cells (Fig. 1 E). In contrast, Patronin::GFP in the ectoderm formed smaller puncta that were distributed across the apical cortex (Fig. 1 B).

Earlier in development, from syncytial to early cellularization stages, Patronin::GFP localized to the two centrosomes above the nucleus (Fig. 1 C). During mid-late cellularization, Patronin::GFP shifted to a more apical localization at cell junctions in both the mesoderm and ectoderm (Fig. 1 C; Video 1), consistent with another study at this developmental stage (Takeda et al., 2018). During gastrulation, medioapical Patronin::GFP foci localized to the center of medioapical myosin patches (Fig. 1 F). This localization mirrors that of active RhoA and ROCK (Mason et al., 2013; Mason et al., 2016). Interestingly, apical constriction of the posterior midgut cells (i.e., the endoderm) also exhibited medioapical Patronin::GFP enrichment (Fig. 1 G). These results demonstrated that medioapical Patronin localization is a fundamental organization shared by apically constricting cells during *Drosophila* gastrulation.

### Medioapical Patronin stabilizes non-centrosomal microtubules

Given the striking and centralized Patronin::GFP localization in mesoderm cells, we next determined if this structure represented a centrosome or another type of microtubule-organizing center. First, we fixed embryos and detected α-Tubulin by immunofluorescence. When viewing cells *en face*, mesoderm cells displayed both punctate and fiber-like structures, suggesting that microtubule are arranged both parallel and perpendicular to the apical surface (Fig. 2, A and B). In contrast, the ectoderm had significantly fewer apical microtubules at the time of mesoderm invagination (Fig. 2 B). Because fixation often destroys dynamic cytoskeletal networks, we next visualized microtubules in live embryos expressing GFP-tagged CLIP-170, a plus-end binding protein (Perez et al., 1999), which gave us the best labeling of microtubules at this stage. During early tissue folding, we observed dense patches of GFP::CLIP170 puncta that co-localized with apical myosin and Patronin (Fig. 2, C and D; Video 2). Thus, these CLIP170 patches that co-localized with Patronin resembled apical microtubule-organizing centers. Consistent with this, at high temporal resolution we observed GFP::CLIP170 comets, which suggested microtubule growth, moving from medioapical patches towards cell edges (Fig. 2, D and E). However, we also observed other directionalities in microtubule growth across the cell apex (Fig. 2 D).

**Figure 2.**
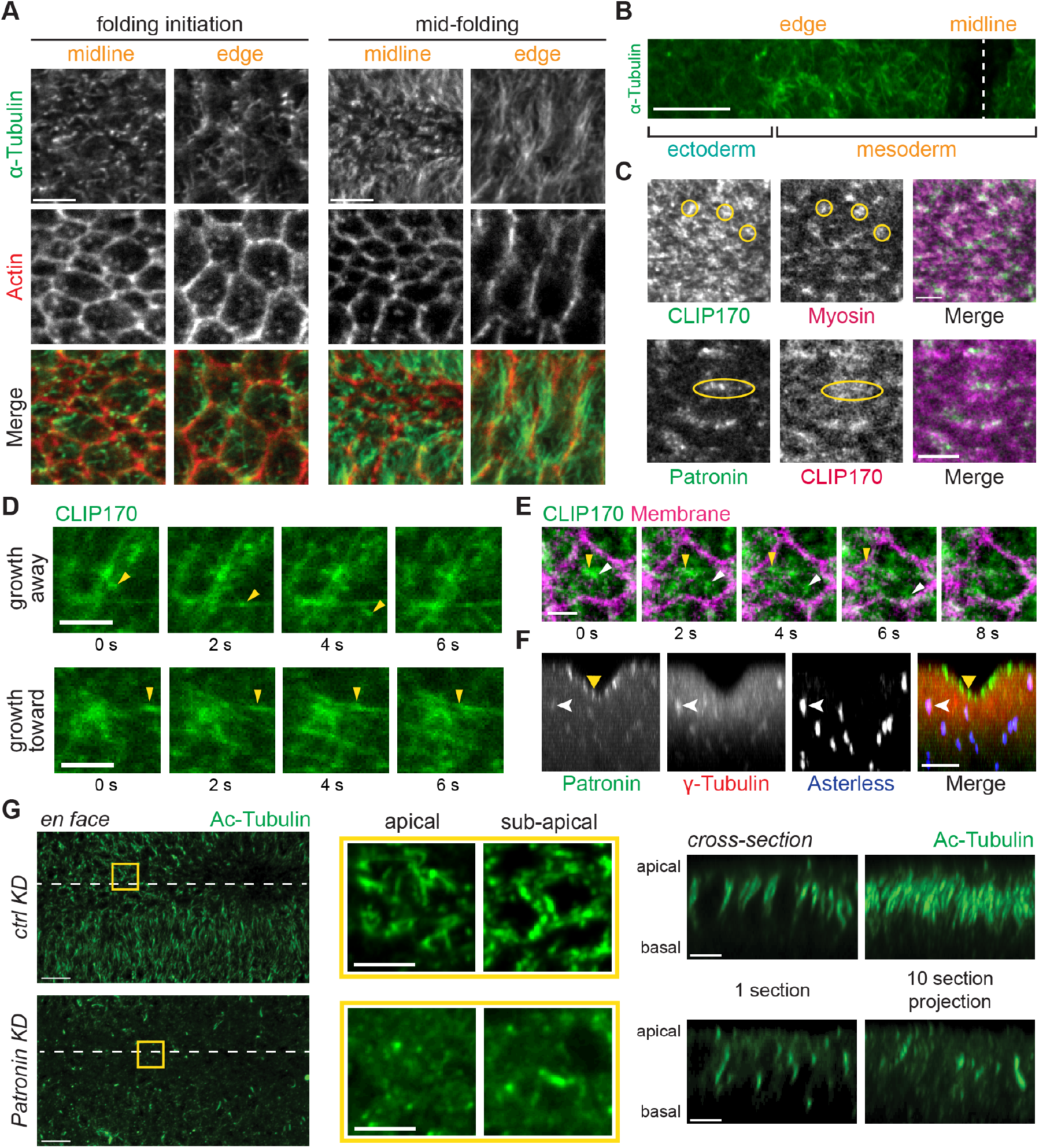
Patronin stabilizes apical, non-centrosomal microtubules in the mesoderm. (A) In the mesoderm, microtubules are oriented parallel and perpendicular to the apical membrane. Images are maximum intensity projections from a fixed embryo stained for α-Tubulin (apical surface) and F-actin (sub-apical section, phalloidin). Surface views of cells along the midline and at the edge of the *snail*-expressing domain are shown for an embryo at folding initiation and mid-folding. (B) Apical microtubules are enriched in mesoderm. Image is an apical surface Z-projection of a fixed embryo stained for α-Tubulin. The furrow midline is marked with a dashed white line. (C) GFP::CLIP170 form dense clusters at the apical surface that co-localize with myosin (yellow circles; top) and Patronin (yellow ovals; bottom). Images are single apical sections from a live movie of an embryo expressing GFP::CLIP170 and Myo::mCH (sqh::mCH) or Patronin::GFP and CH::CLIP170. (D) Microtubule growth can be observed with dynamic GFP::CLIP170 comets (yellow arrowhead). Images are a montage of single apical slices from a live movie of an embryo expressing GFP::CLIP170. (E) Microtubules grow from an apical microtubule-organizing center towards cell edges. Images are a montage of single apical slices from a live movie of an embryo expressing GFP::CLIP170 and Gap43::mCH. Different GFP::CLIP170 comets are marked with arrowheads. (F) Apical Patronin foci do not co-localize with centrosomal markers. Images are maximum intensity projections of apical-basal crosssections from a fixed embryo stained for γ-Tubulin (associated with centrosomes; Dictenberg et al., 1998) (red), Asterless (Cep152, a centriolar component; Varmark et al., 2007; Dzhindzhev et al., 2010) (blue), and endogenous Patronin::GFP signal (green). Medioapical Patronin (yellow arrowhead) is separate from Patronin::GFP signal at centrosomes (white arrowhead). (G) Depleting Patronin destabilizes microtubules. Images are apical surface projections of fixed embryos stained for acetylated-Tubulin. The surface view of Rhodopsin-3 control RNAi (top) and *patronin* RNAi (bottom) are shown on the left. The ventral midline is marked by a white dashed line. Middle images are a magnified *en face* view of apical and sub-apical projections in the yellow box. The right set of images show cross-sections of the same embryos on the left. The left image is a single yz section and the right image is a maximum intensity projection of 10 crosssections. Scale bars, 10 μm (**B; G, tissue-wide view**), 5 μm (**A; C, top; F; G, magnified view and cross-section**), 3 μm (**C, bottom; D-E**).

Apical Patronin is often associated with non-centrosomal microtubules in epithelia (Noordstra et al., 2016; Toya et al., 2016). To test whether these Patronin foci were centrosomes, we examined the position of centrosomal markers, which were significantly below the apical surface throughout folding (Fig. 2 F; Fig. S1A). We observed lower levels of Patronin::GFP associated with centrosomes, which were well-separated from medioapical Patronin::GFP foci (Fig. 2 F). This localization could reflect sites of active Patronin-mediated minus-end stabilization of microtubules that are released from centrosomes, similar to a proposed function of ninein (Mogensen et al., 2000; Moss et al., 2007), or a separate centrosomal pool of Patronin. Thus, there is a distinct organization of non-centrosomal microtubules at the apical cortex during mesoderm invagination.

A well-known function of CAMSAPs is to stabilize microtubules (Goodwin and Vale, 2010; Tanaka et al., 2012; Hendershott and Vale, 2014; Jiang et al., 2014). To determine whether Patronin stabilizes non-centrosomal microtubules near the apical cortex, we fixed Patronin-depleted embryos and immunostained with antibodies against acetylated-α-Tubulin, a marker of stable microtubule polymers (Westermann and Weber, 2003). We verified that RNAi depleted Patronin protein levels by Western blot (Fig. S1 B). Control or wild-type embryos exhibited acetylated microtubules that were both lying across the apical surface and parallel to the apical-basal axis (Fig. 2 G). In contrast to controls, Patronin depletion dramatically reduced visible bundles of acetylated-α-Tubulin (8/8 embryos imaged) (Fig. 2 G). When we examined microtubules in live embryos by imaging GFP::CLIP170, we observed that the organization of apical microtubules was disrupted in Patronin depleted embryos during gastrulation, with centrosomes abnormally localized close to the apical surface (5/5 embryos imaged) (Fig. S1 C).This result suggested that Patronin stabilizes apical, non-centrosomal microtubules and promotes their organization.

Finally, to determine whether Patronin localization required microtubules, we injected Patronin::GFP expressing embryos with the microtubule depolymerizing drug colchicine. Colchicine injection dramatically disrupted Patronin localization (Fig. S1 D), consistent with the reported loss of polarized CAMSAP3 localization in Caco-2 cells treated with nocodazole, another inhibitor of microtubule polymerization (Toya et al., 2016). Together, these results suggested that medioapical Patronin foci form a non-centrosomal microtubule-organizing center in constricting mesoderm cells and that Patronin and microtubule localization is interdependent.

### Patronin puncta coalesce into foci during myosin pulses

During mesoderm invagination, apical myosin initially accumulates in cycles of assembly and disassembly and these pulses are associated with apical constriction (Martin et al., 2009; Vasquez et al., 2014). To determine how medioapical Patronin foci form, we analyzed the spatiotemporal dynamics of Patronin::GFP relative to myosin. In contrast to myosin, we did not observe clear cycles of assembly and disassembly for Patronin::GFP puncta (Fig. 3 A; Video 3). Instead, medioapical Patronin foci appeared to grow by the continuous coalescence of smaller Patronin::GFP puncta (Fig. 3 A). To determine if Patronin::GFP coalescence was associated with apical myosin contraction, we identified individual myosin pulses and analyzed the behavior of Patronin::GFP in the region of the pulse. During myosin pulse assembly, we usually observed instances of Patronin::GFP coalescence, which formed a brighter, more compact focus (Fig. 3 B). When we analyzed the maximum intensity profiles of myosin and Patronin averaged across 20 pulses, we found that myosin intensity peaked about 5 seconds before maximum Patronin::GFP intensity (Fig. 3 C). These data suggested that actomyosin pulses form medioapical Patronin foci, which we showed co-localize with apical myosin patches (Fig. 1 F).

**Figure 3.**
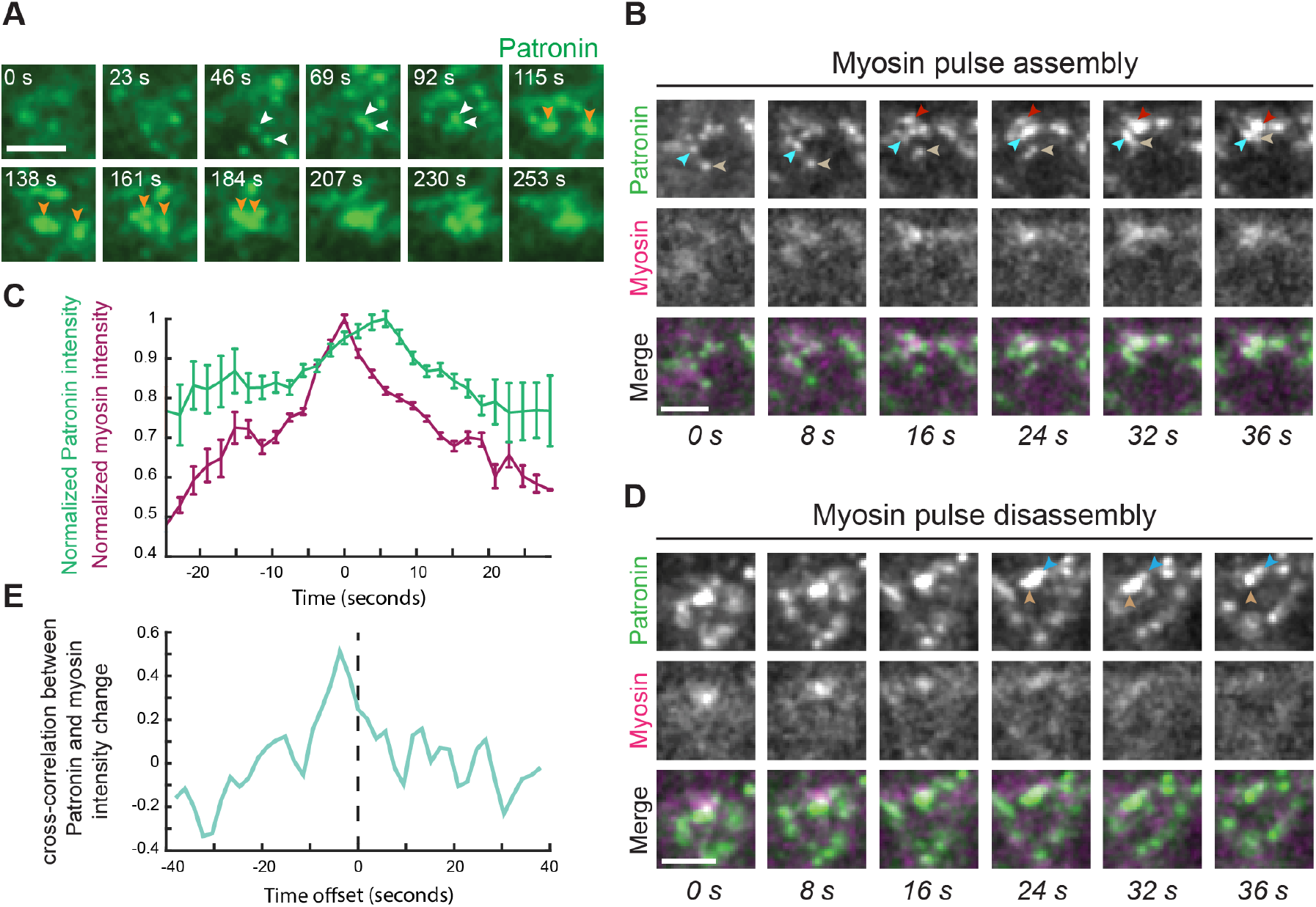
Medioapical Patronin foci form by coalescence during myosin assembly. (A) Patronin foci form by coalescence of smaller puncta (arrowheads). Time-lapse images are from a single apical section from a live movie of an embryo expressing Patronin::GFP. (B) Patronin puncta coalescence (arrowheads) occurs during myosin pulse assembly. Time-lapse images represent a myosin pulse (single apical slice) from a live movie of an embryo expressing Patronin::GFP and Myo::mCH. (C) Quantification of maximum normalized Myo::mCH and Patronin::GFP intensity (n = 4 embryos, 5 pulses per embryo) that has been averaged by aligning the maxima of Myo::mCH across all 20 pulses. Time at peak mean myosin intensity was centered at 0. Error bars represent the standard error of the mean. (D) During myosin pulse disassembly, local Patronin::GFP intensity decreases and larger foci revert back to smaller puncta (arrowheads). Time-lapse images are a representative pulse from a single apical section from a live movie of an embryo expressing Patronin::GFP and Myo::mCH. (E) Myosin and Patronin intensity are tightly correlated. Plotted is the mean time-resolved correlation function between changes in Myo::mCH and Patronin::GFP intensity. Maximum correlation occurs at an offset of ~-4 seconds. Dashed line indicates 0 time offset. Scale bars, 3 μm.

During myosin pulse disassembly, there were local decreases in Patronin::GFP intensity as more compact Patronin foci sometimes appeared to relax and revert back to separate, smaller puncta (Fig. 3, C and D). To determine how closely changes in Patronin intensity tracked with myosin intensity, we analyzed the time-resolved cross-correlation between changes in Patronin::GFP and myosin intensities. We identified a significant correlation that occurred between these two signals, with the maximum correlation occurring at a time offset of −3.9 seconds, consistent with increases in myosin signal preceding increases in Patronin signal (Fig. 3 E). These results demonstrated a tight correlation between myosin and Patronin intensity with myosin preceding Patronin coalescence, suggesting that actomyosin contraction forms medioapical Patronin foci in mesoderm cells, possibly through an advection-based mechanism (Munro et al., 2004; Munjal et al., 2015).

### Actomyosin contraction forms a medioapical, non-centrosomal microtubule-organizing center

Because microtubule/Patronin organization in the mesoderm resembled that of active RhoA and was correlated with myosin contractility (Fig. 3) (Mason et al. 2013; Mason et al., 2016), we tested whether RhoA and myosin contractility were required to organize apical non-centrosomal microtubules. Because ROCK is the main myosin kinase in early *Drosophila* embryos, we injected embryos with the ROCK inhibitor, Y-27632 (Royou et al; 2002; Dawes-Hoang et al., 2005). In comparison to water-injected embryos, Y-27632 disrupted Patronin::GFP apical localization (Fig. 4, A and B). However, because Y-27632 also inhibits aPKC (Davies et al., 2000), we additionally analyzed Patronin localization after inhibiting RhoA activity with the C3-exoenzyme (Crawford et al., 1998). RhoA inhibition similarly disrupted apical Patronin (Fig. 4, C and D), suggesting that RhoA activity promotes the formation of a medioapical microtubule-organizing center.

**Figure 4.**
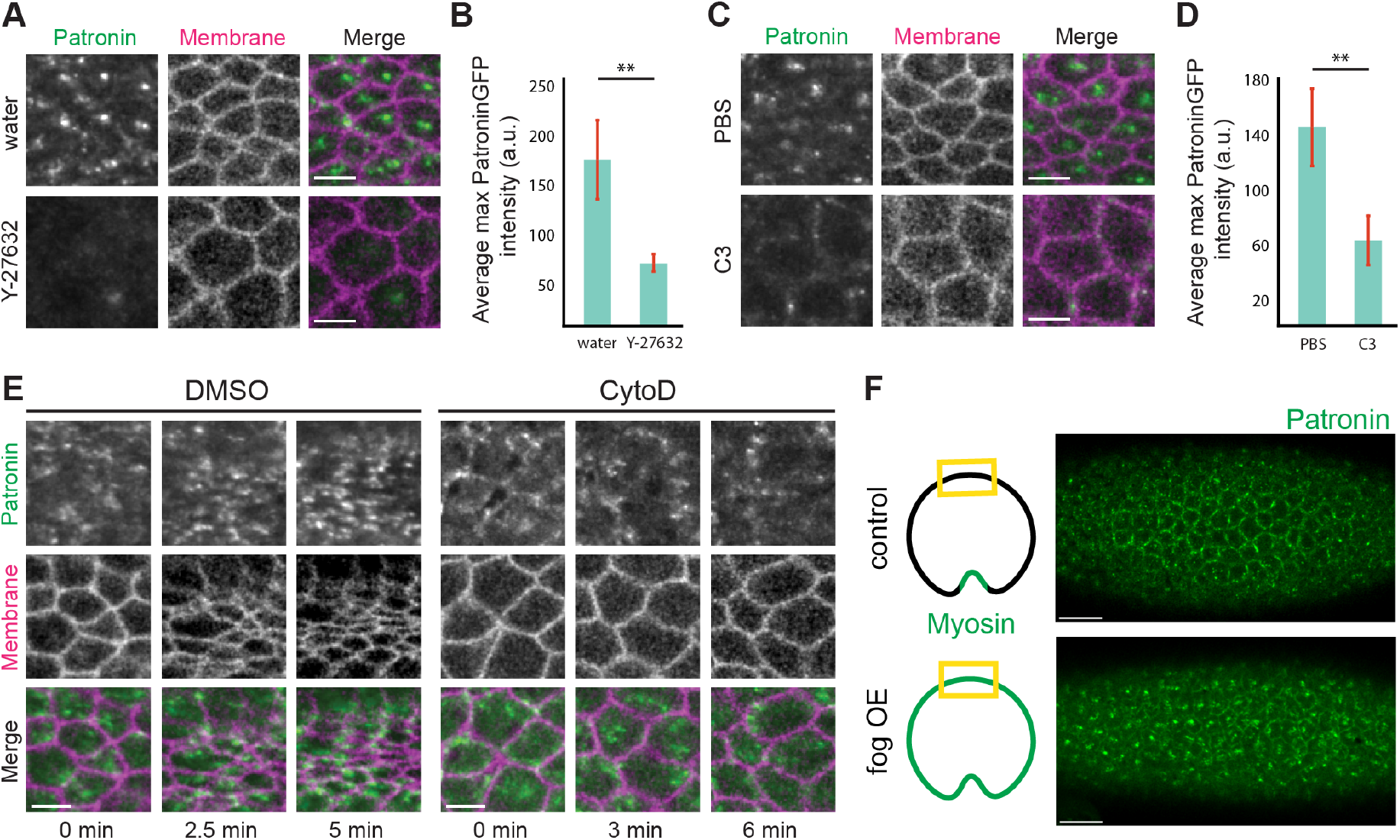
Medioapical Patronin foci depend on actomyosin contraction. (A) ROCK inhibitor disrupts apical Patronin localization in the mesoderm. Images are maximum intensity Z-projections of mesoderm cells from live embryos expressing Patronin::GFP (apical surface) and Gap43::mCH (sub-apical section) injected with water (top) or Y-27632 (50 mM; bottom). (B) ROCK inhibitor decreases Patronin::GFP apical intensity. Quantification of maximum Patronin::GFP intensity in a region that spans the apical cortex (n = 7 measurements per embryo, 4 embryos per condition; **, P < .0001, unpaired t test). Error bars represent one standard deviation. (C) RhoA inhibition disrupts medioapical Patronin localization. Images are maximum intensity Z-projections from a live embryo expressing Patronin::GFP (apical surface) and Gap43::mCH (sub-apical section) injected with PBS (top) or the C3 exoenzyme (1 mg/mL; bottom). (D) RhoA inhibition decreases apical Patronin::GFP intensity. Quantification of maximum Patronin::GFP intensity in a region that spans the apical cortex (n = 7 measurements per embryo, 4 embryos per condition; **, P < .0001, unpaired t test). Error bars represent one standard deviation. (E) Disrupting the apical F-actin network disrupts medioapical Patronin foci. Time-lapse images are maximum intensity Z-projections from live embryos expressing Patronin::GFP (apical surface) and Gap43::mCH (sub-apical section) injected with DMSO (left) and CytoD (.125 mg/mL; right). (F) *Fog* signaling is sufficient to form medioapical Patronin foci in the ectoderm. Images are maximum intensity projections of control and *fog* overexpressed embryos expressing Patronin::GFP. The tissue region shown in the images are highlighted by yellow boxes in the cartoon diagrams. Scale bars, 15 μm (**F**) and 5 μm (**A-E**).

Because RhoA also promotes contraction via F-actin assembly (Evangelista et al., 1997; Watanabe et al., 1997; Großhans et al., 2005; Otomo et al., 2005; Fox and Peifer, 2007; Murrell et al., 2015; Agarwal and Zaidel-Bar, 2018), we tested whether disrupting the apical F-actin network affects microtubule organization by injecting drugs that interfere with F-actin assembly, such as Cytochalasin D (CytoD) and Latrunculin B (LatB). Similar to Y-27632 and C3-exoenzyme injections, both CytoD and LatB disrupted medioapical Patronin foci formation. In these embryos, Patronin puncta did not coalesce and remained as small, dynamic puncta (Fig. 4 E; Fig. S2 A). Importantly, it has been shown in Caco-2 cells, where CAMSAP-3 localizes as puncta in the apical cortex, that F-actin depolymerization reduces CAMSAP-3 puncta in cortical regions (Toya et al., 2016). Many of these smaller Patronin puncta co-localized with myosin, suggesting physical interactions between the two cytoskeletal networks (Fig. S2 B). Altogether, our data indicated that RhoA activity, which leads to actomyosin contraction, is required to organize apical, non-centrosomal microtubules during mesoderm invagination.

RhoA activity at the apical surface in apically constricting cells during *Drosophila* gastrulation is downstream of G-protein-coupled receptor signaling that is activated by the extracellular ligand Folded gastrulation (Fog) (Costa et al., 1994; Dawes-Hoang et al., 2005). Thus, we tested whether *fog* signaling was sufficient for organizing apical, non-centrosomal microtubules. Ectopic *fog* expression outside the mesoderm leads to apical myosin activation across the entire surface of the embryo (Dawes-Hoang et al., 2005). When we looked at Patronin::GFP localization in ectodermal tissues in *fog* expressing embryos, we observed medioapical foci of Patronin::GFP (10/10 embryos imaged), in contrast to the largely junctional localization of Patronin::GFP in control embryos as described above (Fig. 1, B and D; Fig. 4 F). Therefore, our data support the hypothesis that RhoA and actomyosin contractility, downstream of *fog* signaling, drive the formation of apical, non-centrosomal microtubule-organizing centers.

### Microtubules are not required for initiating apical contractility or adherens junction assembly

Studies in other developmental contexts have shown that microtubules are necessary for cell invagination (Lee et al., 2007; Booth et al., 2014). Consistent with this, we found that microtubules were also important for mesoderm cell invagination (Fig. 5 A; Video 4). Given the striking organization of the microtubule cytoskeleton in the mesoderm, we sought to determine how microtubules promote cell invagination. For example, microtubules could help induce contractility, such as by activating apical actomyosin assembly (Rogers et al., 2004; Booth et al., 2014). Another possibility is that microtubules could regulate adherens junction assembly or position to promote cell adhesion and/or force transmission (Harris and Peifer, 2005; Stehbens et al., 2006; Meng et al., 2008; Le Droguen et al., 2015). Furthermore, microtubules might regulate the connection between actomyosin networks and the adherens junctions.

**Figure 5.**
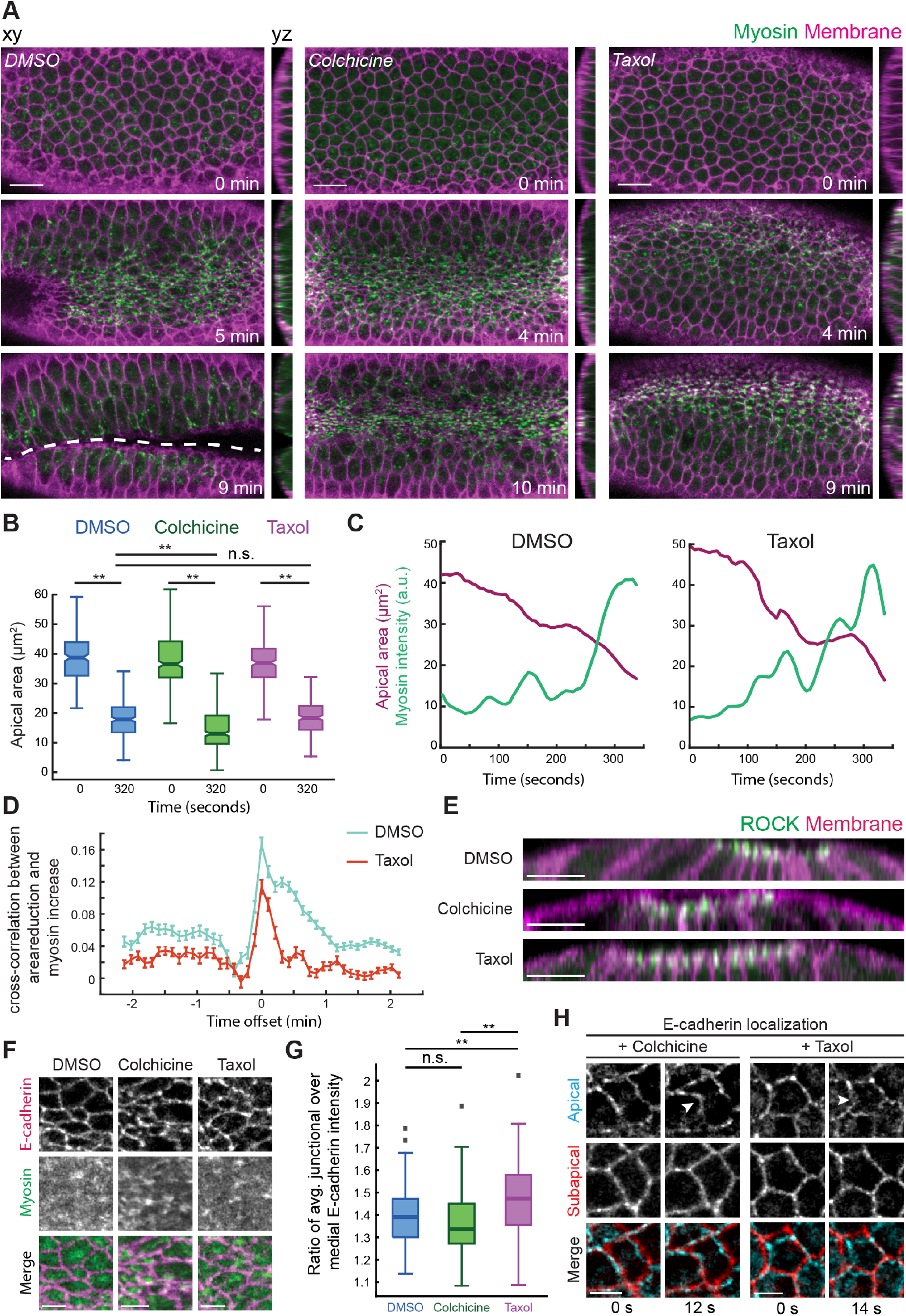
Apical myosin activation and apical constriction initiation do not require microtubules. (A) Disrupting microtubules disrupts folding, despite apical myosin activation and apical constriction initiation. Time-lapse images are maximum intensity Z-projections from live embryos expressing sqh::GFP (myosin, apical surface) and Gap43::mCH (membrane, sub-apical section) injected with DMSO (left), colchicine (5 mg/mL; middle), and taxol (5 mg/mL; right). Apical-basal cross-sections (yz) are to the right of each image. Dashed line indicates the ventral furrow. (B) Apical area initiates reduction after disrupting microtubules. Quantification of apical cell areas preconstriction (t = 0 s) and 320 s after from three representative live embryos injected with DMSO (n = 226 cells, T = 0 s; 252 cells, T = 320 s; **, P < .0001, unpaired t test), colchicine (n = 284 cells, T = 0 s; 353 cells, T = 320 s; **,P < .0001, unpaired t test), and taxol (n = 275 cells, T = 0 s; 226 cells, T = 320 s; **,P < .0001, unpaired t test). Notch is the median, bottom and top edges of the box are the 25^th^ and 75^th^ percentiles, whiskers extend to the most extreme data points. (C) Myosin assembly and pulses still occur after inhibiting microtubule dynamics. Graphs show apical area and myosin intensity over time in representative single cells in DMSO- (top) and taxol-injected (bottom) embryos. (D) Disrupting microtubule dynamics does not initially interfere with the correlation between myosin increase and area reduction. Plotted is the mean time-resolved correlation function between constriction rate (+ constriction = area decrease) and myosin rate. Data are from 3 representative embryos injected with DMSO (n = 548 cells) or taxol (n = 543 cells). Error bars represent the standard error of the mean. (E) Microtubule disruption does not affect apical ROCK polarity. Images show apical-basal cross-sections from representative live embryos expressing ROCK::GFP and Gap43mCH and injected with DMSO, colchicine, or taxol. (F) Apical spot adherens junctions are unaffected by microtubules disruption. Images are apical surface projections of a live embryo expressing E-cadherin::GFP and Myo::mCH injected with DMSO (left), colchicine (middle), and taxol (right). (G) Quantification of junctional E-cadherin enrichment. Individual cells at the onset of constriction were segmented and the ratio of average raw junctional and medioapical Ecadherin::GFP intensity was calculated for 2-3 representative DMSO- (n = 122 cells), colchicine- (n = 67 cells), and taxol-injected (n = 105) embryos. (**, P <.0001, unpaired t test). Bottom and top edges of the box are the 25^th^ and 75^th^ percentiles, whiskers extend to the most extreme data points; outliers are included (gray squares). (H) Apical spot junctions are still pulled in after microtubule disruption. Time-lapse images showing a single apical (cyan) and sub-apical (red) slice of representative embryos expressing E-cadherin::GFP injected with colchicine (left) and taxol (right). White arrowheads point to an apical spot junction that has been pulled inwards in the second frame. Scale bars, 15 μm (**A**), 10 μm (**E**) and 5 μm (**F-H**).

To determine whether microtubules are required to initiate contractility in mesoderm cells, we disrupted microtubules pharmacologically or by gene depletion. Embryos staged at late cellularization were acutely injected with drugs that disrupt the microtubule cytoskeleton and were imaged within minutes post injection. Despite tissue folding failure, we found that colchicine injection did not disrupt apical myosin activation or the initiation of apical constriction (19/19 embryos imaged) (Fig. 5 A; Video 4). To demonstrate that apical constriction occurred, we measured apical area at the onset of constriction and ~ 5 minutes later for both control and drug-injected embryos (Fig. 5 B). Depleting Patronin by RNAi also disrupted tissue folding (11/44 embryos imaged) without affecting mesoderm cell fate, apical myosin localization, or constriction onset (44/44 embryos imaged) (Fig. S3, A - C). Interestingly, Patronin depletion caused an initial heterogeneity in apical cell area (44/44 embryos imaged), consistent with prior observations in *Drosophila* embryos (Takeda et al. 2017)

To test whether microtubule dynamics/organization were important for apical force generation, we injected embryos with taxol, a microtubule-stabilizing agent. Taxol injection disrupted apical microtubule organization, resulting in thick bundles that spanned the apical surface (Fig. S3 D). Perturbing microtubule dynamics/organization with taxol disrupted folding, but again, had no effect on initial cell constrictions or myosin accumulation (21/21 embryos imaged) (Fig. 5, B and C). Furthermore, taxol-injected embryos exhibited a clear cross-correlation peak between apical constriction rate and the rate of change in myosin intensity, which is indicative of normal myosin pulsing (Fig. 5 D) (Mason et al., 2013; Vasquez et al., 2014). Finally, the apically polarized localization of ROCK was unaffected in both colchicine- and taxol-injected embryos (Fig. 5 E). These data suggested that microtubule organization is not important for apical actomyosin activation or the subsequent onset of apical constriction.

To determine if microtubules were required for apical adherens junction assembly (Dawes-Hoang et al., 2005; Stehbens et al., 2006; Kölsch et al., 2007; Marston et al., 2016; Weng and Wieschaus, 2016), we analyzed E-cadherin in live embryos injected with microtubule drugs or depleted of Patronin. There were no gross defects in apical E-cadherin structure or polarity in either case (Fig. 5 F; Fig. S3, B and C). We measured raw intensity values of E-cadherin::GFP at the onset of constriction and calculated the average ratio of junctional over medial E-cadherin intensity in DMSO-, colchicine-, and taxol-injected embryos. In all cases, E-cadherin was present and junctionally enriched (Fig. 5 G). Consistent with proper adherens junction assembly, we observed wavy and deformed cell-cell interfaces at the apical surface, which suggested that medial actomyosin was initially able to pull on spot adherens junctions (Fig. 5 H). Thus, during mesoderm invagination, microtubules are not necessary to initiate contractility (i.e., apical myosin activation) or to assemble apical adherens junctions, but are still required for tissue folding.

### Proper microtubule organization promotes intercellular force transmission

In the previous section, we showed that initiation of apical myosin and adherens junction accumulation was largely unaffected in Patronin-depleted, colchicine-, and taxol-injected embryos. Another possibility was that microtubules promote the connection between actomyosin and adherens junctions and/or promote reattachment of lost connections (Roh-Johnson et al., 2012; Jodoin et al., 2015). Therefore, we analyzed actomyosin attachments to adherens junctions and the resulting intercellular connectivity between actomyosin networks after microtubule cytoskeleton disruption. In both colchicine- and taxol-injected embryos, the apical myosin networks exhibited striking separations well after apical constriction onset (Fig. 6, A and B; Fig. S4 A; Video 5). Myosin spots in different cells chaotically separated and then moved back together and the apex of cells became distended, losing their constricted morphology (Fig. 6 B; Fig. S4 A; Video 5). The loss of intercellular connectivity of actomyosin after microtubule organization disruption was not associated with a fracture of the medioapical ROCK/myosin signaling center, because embryos expressing GFP-tagged ROCK injected with taxol exhibited ROCK/myosin foci that clearly separated from the junctional domain (Fig. S4 B). The phenotypes were not caused by disrupting a previous developmental process, such as cellularization, because injecting colchicine into live embryos that had completed cellularization and started apical constriction (i.e., injected when apical myosin was present) resulted in the same phenotype (2/2 embryos imaged) (Fig. S4 C). Thus, microtubules are required during folding to maintain actomyosin attachments to adherens junctions.

**Figure 6.**
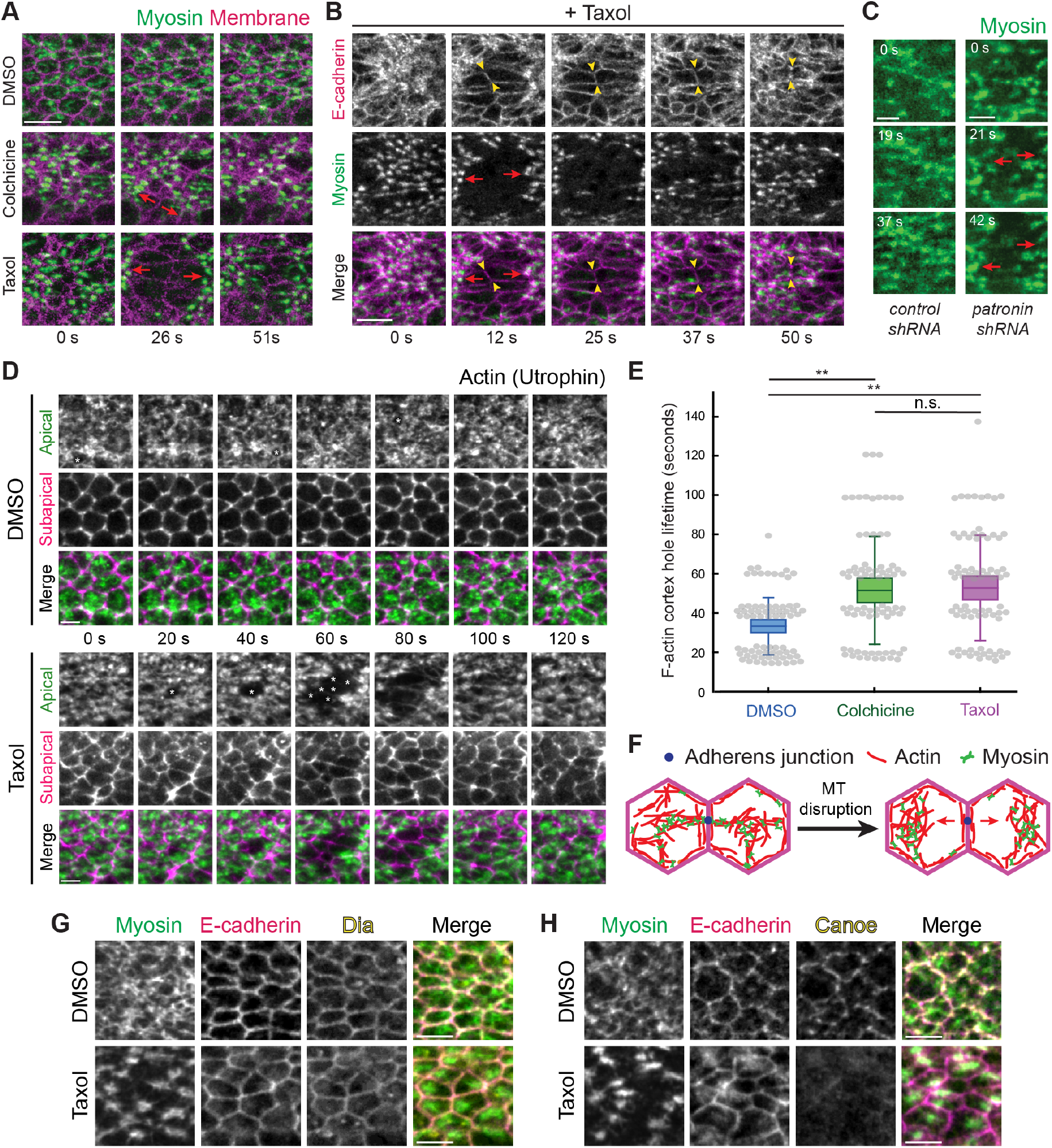
Microtubules stabilize actomyosin connections to intercellular junctions. (A) Microtubule disruption leads to separations between myosin structures and intercellular junctions. Time-lapse images are maximum intensity Z-projections from live embryos expressing Myo::GFP (apical surface) and Gap43::mCH (sub-apical section illustrating junctions) injected with DMSO (top), colchicine (5 mg/mL; middle), and taxol (5 mg/mL; bottom). Red arrows indicate the direction in which myosin structures separate from cell junctions. (B) E-cadherin is still present at interfaces between myosin spot separations. Time-lapse images are maximum intensity Z-projections of apical Myo::mCH and an apical Z-slice of E-cadherin::GFP for a representative embryo injected with taxol (5 mg/mL). Red arrows indicate the direction in which myosin spots separate from cell junctions. One cell-cell interface is highlighted between yellow arrowheads. (C) Depleting Patronin also causes myosin network separation. Time-lapse images are apical projections from representative live embryos expressing Myo::GFP and shRNA against Rhodopsin-3 (control, top) and Patronin (bottom). Red arrows indicate the direction in which myosin spots separate from cell junctions. (D) Taxol injection causes longer-lived and larger holes in the F-actin cortex (asterisks) which lead to separation of F-actin meshworks away from junctions. Time-lapse images are maximum intensity Z-projections of apical slices (green) and a subapical slice (magenta) from representative live embryos expressing Utr::GFP injected with DMSO or taxol (5 mg/mL). (E) The lifetime of holes in the F-actin cortex near cell junctions is longer when microtubules are disrupted. Quantification of hole lifetime length at a resolution of ~20 seconds between time steps for 75 holes across 3 embryos in each condition (**, P <.0001, unpaired t test). Bottom and top edges of the box are the 25^th^ and 75^th^ percentiles, whiskers extend to the most extreme data points, excluding outliers. All data points were plotted as gray dots. (F) Diagram showing model for myosin separation. Shown is a top down view of the apical cortex in two adjacent cells. Loss of attachment of actomyosin to the adherens junction (blue dot) after microtubule disruption leads to separation. (G) Microtubule disruption does not affect junctional localization of Dia. Images are from embryos expressing Myo::GFP that were injected with DMSO (left) or taxol (5 mg/mL; right), PFA fixed, and immunostained with antibodies against E-cadherin and Dia. (H) Microtubule disruption affects junctional localization of Cno. Images are from embryos expressing Myo::GFP that were injected with DMSO (left) or taxol (5 mg/mL; right), PFA fixed, and immunostained with antibodies against E-cadherin and Cno. Scale bars, 10 μm (**A-B**), 5 μm (**C-H**).

When we perturbed microtubules by depleting Patronin by RNAi, which we showed destabilized apical non-centrosomal microtubules and disrupted their organization (Fig. 2 G; Fig. S1 C), we also observed dynamic tearing of the myosin network in 11 out of 44 knockdowns (Fig. 6 C). We believe that the lower penetrance of the *patronin*-RNAi treatment reflects the less severe effect of Patronin depletion on microtubule organization. In both drug-injected and Patronin-depleted embryos, these separations only occurred after a significant buildup of apical myosin and the initial formation of a supracellular myosin network, suggesting that microtubules are important at later stages of the folding process when cells must stabilize their shape in the face of significant tension.

To determine how actomyosin network structures separated from each other, we examined the F-actin cortex using a GFP-tagged F-actin-binding domain of Utrophin (Utr::GFP) (Rauzi et al., 2010). Similar to wild-type embryos (Jodoin et al., 2015), DMSO-injected embryos exhibited a dynamic F-actin cortex where apical F-actin holes appeared next to intercellular junctions and were rapidly repaired, usually within ~20 - 60 seconds (Fig. 6, D and E). In contrast, taxol-injected embryos exhibited longer-lived holes or fractures in the F-actin meshwork and these fractures often grew larger to encompass multiple neighboring cells (Fig. 6, D and E; Video 6). These results suggested that actomyosin network separations occur due to the separation of the apical F-actin meshwork from intercellular junctions (Fig. 6, D - F). We previously showed that E-cadherin was still present at junctions after microtubule disruption (Fig. 5 F; Fig. S3, B and C). To determine whether the actomyosin cortex separates from adherens junctions, we visualized E-cadherin and found that E-cadherin persists at the cell-cell interface during myosin separations (Fig. 6 B). Moreover, actomyosin network separations were repaired as myosin in neighboring cells re-established connections and pulled back together (Fig. 6 B; Fig. S4 A; Video 5), which further suggested the presence of functional adherens junctions. Only at much later stages, when the tissue integrity was completely lost, did we observe E-cadherin mis-localized across the apical cortex (Fig. S4 D).

Because actin turnover reattaches the F-actin cortex to the junctions (Jodoin et al.), the greater persistence of the F-actin holes after microtubule disruption suggested that microtubules could promote force transmission by inducing actin turnover. We examined whether microtubules influenced the localization of known F-actin binding proteins that localize to junctions. We fixed embryos after either DMSO or taxol injection and stained for the formin Diaphanous (Dia), which is enriched at junctions (Mason et al., 2013), and Canoe (Cno), the *Drosophila* Afadin homologue which mediates linkages between F-actin and adherens junctions (Sawyer et al., 2009; Choi et al., 2016). In both conditions, we observed a clear junctional enrichment of Dia (5/5 DMSO embryos; 6/6 taxol embryos) (Fig. 6 G). However, we cannot rule out that microtubules promote Dia activity, as CLIP170 has been shown to stimulate mDia1 activity (Henty-Ridilla et al., 2016). In contrast, we observed a striking loss of junctional Cno localization after taxol injection (4/4 DMSO embryos; 4/4 taxol embryos) (Fig. 6 H). These results suggested that proper microtubule organization is required for Cno localization at adherens junctions, which could promote F-actin recruitment and linkage to the adherens junction (see Discussion). Overall, our results demonstrated that microtubules are critical for actomyosin networks in adjacent cells to stably transmit force to each other across adherens junctions by promoting reattachment of apical actin meshwork to the junctions (Fig. 6 F).

## Discussion

Our work identifies a role for microtubules in promoting force transmission between epithelial cells during *Drosophila* mesoderm invagination. We have demonstrated, to our knowledge, a novel organization to the microtubule cytoskeleton, where a medioapical focus of the microtubule minus-end-binding protein Patronin is present in apically constricting cells. Patronin puncta coalesce to form medioapical Patronin foci, which depend on actomyosin contraction. Patronin stabilizes and organizes apical, non-centrosomal microtubules, which grow from this center to intercellular junctions. Microtubules are dispensable for apical myosin polarity, apical E-cadherin, and initiation of apical constriction, but are required to promote medioapical actomyosin network attachment to E-cadherin-based cell junctions. Disrupting microtubules resulted in dynamic separations between myosin and intercellular junctions that prevented mesoderm invagination. This study uncovers a previously unrecognized role for microtubule organization in integrating contractile forces across a tissue during morphogenesis, possibly by regulating actin turnover.

### Apical constriction in the early *Drosophila* embryo is associated with a medioapical, non-centrosomal microtubule-organizing center

Apically constricting cells in the mesoderm and endoderm shared a similarly polarized microtubule organization with Patronin foci localized to the medioapical cortex. In addition, live imaging of embryos expressing GFP::CLIP170 revealed the appearance of dense patches of microtubules in the mesoderm that co-localized with both apical myosin patches and Patronin foci. Microtubules were observed to grow from medioapical patches towards cell junctions. Given the high concentration of microtubule-associated proteins that localize to the medioapical focus, this resembles a type of non-centrosomal microtubule-organizing center. It is unclear how similar this non-centrosomal microtubule organization is to the organization that has been observed for centrosomal microtubules in MCF-7, MDCK, and hE-CHO cells (Stehbens et al., 2006), where microtubules radiate out from centrosomes with growing plus ends enriched towards cell edges. However, in support of this view, both fixed and live data suggested that microtubules span the apical cortex and we observed CLIP170 puncta at cell junctions. Together, these results suggest that the medioapical cortex is a zone of microtubule minus-end stabilization that could act as an organizing center from which microtubules grow towards cell junctions (Fig. 7 A).

**Figure 7.**
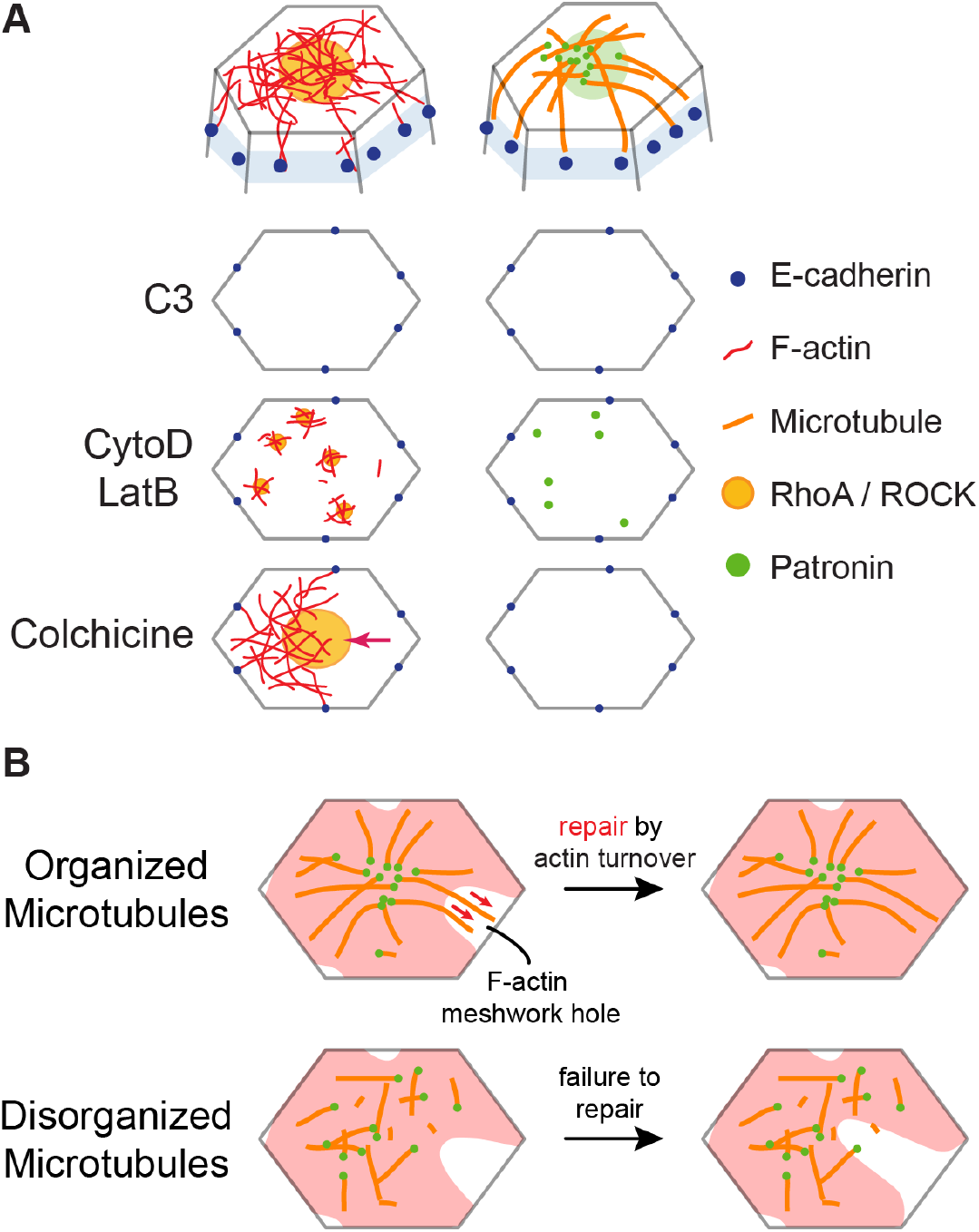
Actin and microtubule cytoskeletons interact to promote intercellular force transmission. (A) Diagrams at top show the proposed organization of the microtubule and actin cytoskeletal networks at the cell apex. Diagrams below show top down views of the apical cortex and the effect of various perturbations on Rho/ROCK/F-actin and Patronin localization. C3 injection eliminates apical RhoA/ROCK and Patronin foci. CytoD/LatB injections lead to formation of smaller RhoA/ROCK and Patronin puncta. Colchicine injection does not affect RhoA/ROCK polarity but leads to actomyosin network separations from intercellular junctions (red arrow). (B) A model for how microtubules promote reattachment of the apical F-actin meshwork to adherens junctions after fracture. In wild-type embryos (top), the organization of apical, non-centrosomal microtubules could promote repair of holes in the F-actin meshwork (red) by guiding F-actin polymerization and/or bundling through physical associations (red arrows). When microtubule organization is disrupted (bottom), the hole in the F-actin meshwork cannot be repaired in a timely manner, leading to actomyosin network separation from adherens junctions.

Patronin foci co-localized with apical myosin patches and formed by coalescence that spatiotemporally correlated with myosin pulses. During myosin pulses, peak Patronin intensity was observed ~4 seconds after peak myosin intensity. Thus, polarized Patronin organization may be a consequence of medioapical actomyosin contraction. Consistent with this, *fog* overexpression was sufficient to reorganize microtubules in ectopic tissues. Furthermore, injecting higher doses of CytoD/LatB, which leads to the formation of small myosin and ROCK puncta that fail to coalesce (Mason et al., 2013; Coravos et al., 2016), caused Patronin to remain as small puncta that do not coalesce (Fig. 7 A). One possible connection between Patronin and F-actin is the spectrin cytoskeleton because spectrin associates with F-actin and Patronin has been shown to associate with spectrin isoforms (King et al., 2014; Khanal et al., 2016). However, Patronin’s central localization also depends on microtubules and it is possible that microtubules trapped in the actin network are ‘collected’ by actomyosin contractile flow through advection (Salmon et al., 2002; Munro et al., 2004; Munjal et al., 2015). To our knowledge, this is the first example of actomyosin contraction forming a microtubule-organizing center.

Additionally, separate from the microtubules that were localized at the apical cortex, we also observed clear arrays and bundles of microtubules along the apical-basal axis (Harris and Peifer, 2007). The interplay between apical-basal microtubule bundles near cell boundaries and the microtubule network at the apical cortex is still unclear. These may integrate with apical, non-centrosomal microtubules or behave as a separate microtubule population that has distinct functions during cell shape change and tissue invagination.

### Microtubules are not required to initiate apical constriction in mesoderm cells

Apical constriction and subsequent tissue invagination have been shown to depend on microtubules in bottle cells during *Xenopus laevis* gastrulation and placodal cells during salivary gland tubulogenesis in *Drosophila* (Lee and Harland, 2007; Booth et al., 2014). In the salivary gland cells, microtubules were required for the formation of a medioapical actomyosin network that constricted the cell apex (Booth et al., 2014), consistent with a proposed model for microtubule-dependent activation of RhoA at the apical cortex (Rogers et al., 2004). Apical cortex organization leading to polarized domains of active contractility is important during apical constriction (Mason et al., 2013; Booth et al., 2014; Coravos et al., 2016). However, in mesoderm cells, depolymerizing microtubules or disrupting microtubule dynamics/organization did not lead to a loss of apical myosin activation and initiation of apical constriction was normal. Moreover, medioapical ROCK polarity was unaffected after taxol injection. Overall, these data suggest that during mesoderm invagination in the early embryo, medioapical RhoA activity is not affected by microtubule perturbation. However, RhoA activity was important for microtubule organization (Fig. 7 A). Our work suggests that microtubules can affect apical constriction in other ways, such as by promoting actin network reattachment to the adherens junction after it fractures.

### Microtubules promote actomyosin reattachment to adherens junctions

The earliest gastrulation phenotype we observed after disrupting microtubules was fracture occurring between apical actomyosin networks and adherens junctions, which failed to be repaired in a timely manner (Fig. 7 B). These apical actin cortex fractures led to the separation of myosin structures and persistent holes in the apical F-actin cortex, which were similar to phenotypes observed in embryos with disrupted actin turnover (Jodoin et al., 2015). Apical microtubules could promote adherens junction reattachment by promoting actin assembly around the adherens junction (Henty-Ridilla et al., 2016), or by promoting F-actin bundling around the adherens junction (López et al., 2014; Michael et al., 2016). These interactions may play a role in repairing F-actin meshwork holes to promote reattachment (Jodoin et al., 2015) (Fig. 7 B). In addition, microtubules may promote actomyosin attachment to adherens junctions by physically associating with the apical F-actin network. Such an association could be mediated by actin-microtubule crosslinkers (Applewhite et al., 2010; Girdler et al., 2016; Takács et al., 2017). When we injected embryos with CytoD or LatB, Patronin localized as small, dynamic puncta across the apical surface of cells, similar to myosin (Mason et al., 2013). Many of these Patronin puncta co-localized with myosin, suggesting a physical association between apical actomyosin and microtubules. While the importance of microtubules for apical constriction initiation could be tissue-specific, it will be important to investigate if microtubules promote actomyosin connections to intercellular junctions in other developmental contexts.

We conclude that microtubules have a specific role for promoting actomyosin attachments to cell junctions for several reasons. The phenotype we observe is not consistent with a severe loss of E-cadherin based cell junctions because: 1) E-cadherin is still present apically at junctions, and 2) actomyosin is able to engage with junctions and reattach after separation, in contrast to mutant embryos with depleted junctions (Martin et al., 2010). While Dia was able to localize to junctions after microtubule disruption, we observed a depletion of Cno at junctions. Loss of Cno leads to a separation of actomyosin from junctions, but prevents reattachment (Sawyer et al., 2009). Because actomyosin was able to reattach, loss of Cno localization may be a consequence of repeated separations of actomyosin from adherens junctions.

## Materials and Methods

### Fly stocks and genetics

Fly stocks and crosses used in this study are listed in Table S1. Control (Rhodopsin 3) and Patronin knockdown flies were generated by crossing virgins of the shRNA lines to male flies carrying maternally-loaded Gal4 drivers with appropriate markers. Crosses were maintained at 27 °C. In the F2 generation, non-balancer females and males were used to set up cages that were incubated at 25 °C. All other crosses and cages were maintained at 25 °C.

### Live and fixed imaging

For live imaging, embryos were dechorionated in 50% bleach, washed in water, and mounted onto a glass slide coated with glue (double-sided tape dissolved in heptane). Coverslips (No. 1.5) coated in glue were attached to the slide to use as spacers and a No. 1 coverslip was attached on top to create a chamber. Halocarbon 27 oil was used to fill the chamber. All imaging took place at room temperature (~ 23 °C).

For fixed imaging, embryos were dechorionated in bleach, washed in water, then fixed in 8% paraformaldehyde in 0.1 M phosphate buffer at pH 7.4 with 50% heptane for 30 minutes and manually devitellinized. For the best microtubule staining, embryos fixed in PFA were devitellinized by removing fixative, adding 50% methanol, and vortexing. These embryos were stored in 100% methanol at −20 °C and rehydrated in .01 % Tween 20 in PBS (PBS-T). In addition, some embryos (Fig. S3 B) were heat-fixed (HF) in boiled Triton salt solution (.03% Triton X-100 and 0.4% NaCl in water), cooled on ice, and devitellinized in a 1:1 heptane/methanol solution and stored and rehydrated as above.

Embryos were washed in PBS-T, blocked with 10% BSA in PBS-T, and incubated with antibodies diluted in PBS-T. To visualize F-actin, manually devitellinized embryos were incubated with Alexa Fluor 647 conjugated phalloidin (Invitrogen) diluted in 5% BSA in PBS-T overnight at 4 °C. For PFA-fixed and manually devitellinized embryos, Asterless was recognized using antibody (a gift from J. Raff) diluted at 1:500, γ-Tubulin (Sigma-Aldrich, GTU-88) at 1:500, Dia (a gift from S. Wasserman) at 1:5000, and Cno (a gift from M. Peifer) at 1:500. For PFA-fixed embryos devitellinized by methanol addition and vortexing, α-Tubulin was recognized using antibody (Sigma-Aldrich) diluted at 1:500 and acetylated-tubulin antibody (Sigma-Aldrich, 6-11B-1) at 1:500. For HF embryos, Armadillo was recognized using antibody (Developmental Studies Hybridoma Bank) diluted 1:500 and Snail using antibody (a gift from M. Biggin) at 1:500. Embryos were incubated with primary antibodies at room temperature for 2 hours. Secondary antibodies used were Alexa Fluor 488, 568, or 647 (Invitrogen) diluted at 1:500 in 5% BSA in PBS-T incubated overnight at 4 °C. Endogenous GFP signal was visualized for Patronin::GFP. After antibody incubation, embryos were mounted onto glass slides using AquaPolymount (Polysciences, Inc.).

All images were taken on a Zeiss LSM 710 confocal microscope with a 40x/1.2 Apochromat water objective lens, an argon ion, 561 nm diode, 594 nm HeNe, 633 HeNe lasers, and Zen software. Pinhole settings ranged from 1 – 2.5 airy units. For two-color live imaging, band-pass filters were set at ~490 – 565 nm for GFP and ~590 – 690 nm for mCH. For three-color imaging, band-pass filters were set at ~480 – 560 nm for GFP, ~580 – 635 nm for Alexa Fluor 568, and ~660 – 750 nm for Alexa Fluor 647.

### Image processing and analysis

All images were processed using MATLAB (MathWorks) and FIJI (http://fiji.sc/wiki/index.php/Fiji). A Gaussian smoothing filter (kernel = 1 pixel) was applied. Apical projections are maximum intensity Z-projections of multiple z sections (2-4 μm) and sub-apical sections are optical slices that are 1 - 2 μm below the apical sections.

Image segmentation for quantifications of cell area and myosin and Patronin intensities was performed using custom MATLAB software titled EDGE (Embryo Development Geometry Explorer; https://github.com/mgelbart/embryo-development-geometry-explorer; Gelbart et al., 2012). To calculate the junctional to medial ratio of Patronin::GFP intensity (Fig. 1 D), raw images were processed via background subtraction to remove cytoplasmic Patronin::GFP by subtracting the mean plus one half standard deviation intensity value for the first time step from every pixel in all images. We made maximum intensity projections via FIJI and imported the image stacks into EDGE. Cell boundaries were automatically detected and manually corrected, after which EDGE exported cell area and integrated intensity of Patronin::GFP for each cell. Medial Patronin::GFP intensity (I_M,Int_) was defined as the integrated intensity value for the whole cell (I_W,Int_) excluding the outermost 2 layers of pixels. Junctional Patronin::GFP intensity (I_J,Int_) was calculated as the difference between I_W,Int_ and I_M,Int_. EDGE exported area values as total pixel numbers comprising the region of interest for both the whole cell (S_W_) and the medial region (S_M_). The area of the junctional region (S_J_) was defined as the difference between S_W_ and S_M_. Then we proceeded to calculate the average pixel intensity of the junctional region (i_J_) to that of the medial region (i_M_) as follows:

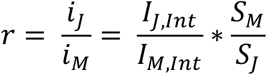

We used average per pixel intensity values instead of the integrated intensity values for that region to more accurately represent the local protein concentration. The same analysis was applied to calculate the junctional to medial ratio of E-cadherin::GFP intensity (Fig. 5 G).

To quantify changes in cell area and myosin intensity (Fig. 5, C and D), raw images were processed via background subtraction as described above. We made maximum intensity projections via FIJI and imported the image stacks into EDGE. Cell boundaries were automatically detected and manually corrected, after which EDGE exported cell area and integrated intensity of Myosin::GFP for each cell. To calculate the cross-correlation between the rate of change in cell area constriction and per pixel myosin intensity buildup rate (Fig. 5 D), we first smoothed the area and myosin integrated intensity curves of each cell by a moving average (3 time steps wide) and then used these values to calculate the average per pixel intensity through time. Then, we found the rate of change in cell area constriction and per pixel myosin intensity buildup rate by finding the difference between a value at a given time step and the value 2 time steps prior, then dividing by the time difference and including only the timepoints between constriction initiation and mesoderm invagination. Finally, we used the MATLAB ‘xcorr’ function to calculate the normalized cross-correlation between these two rates. We aggregated cross-correlation curves from multiple cells across three embryos from each condition (DMSO- or taxol-injected) and obtained the average cross-correlation plot.

For the myosin and Patronin pulse analysis, we manually identified 5 myosin pulses in each of the 4 embryos via FIJI and drew an elliptical ROI around each pulse. We recorded the maximum pixel intensity in manually identified ROIs for both. To plot the average relative behavior of the Myo::mCherry and Patronin::GFP signals (Fig. 3 C), we smoothed the intensity data for each pulse by a moving average (3 time steps wide) and aligned them such that peak myosin intensity for all pulses would fall at a relative time offset of 0 seconds. Then, we calculated average Myo::mCherry and Patronin::GFP intensities for each relative time offset. We found the normalized crosscorrelation between the rates of change in myosin intensity and Patronin intensity (Fig. 3 E) as indicated above.

### Immunoblotting

Early gastrula stage embryos were collected and homogenized in SDS sample buffer, boiled for 5 minutes, and kept on ice. Samples were run on Mini-PROTEAN TGX Precast Gels (Bio-Rad). Primary antibodies used for immunoblotting included α-Tubulin (1:500; Sigma-Aldrich) and Patronin (antibody serum; 1:50; a gift from Ron Vale). Primary antibodies were detected by horseradish peroxidase-labeled secondary antibodies. Signal was developed using SuperSignal West Femto Maximum Sensitiity Substrate (ThermoFisher).

### Drug injections

Dechorionated embryos were mounted onto glass slides and desiccated for 4 minutes using Drierite (Drierite). Embryos were covered with a 3:1 mixture of halocarbon 700/halocarbon 27 oils and then injected laterally during mid-late cellularization. For ROCK inhibition, Y-27632 was dissolved in water and injected at 50 mM concentration. Seventeen embryos were imaged and quantifications in Fig. 4 B are from 4 representative embryos. As a control, water was injected. For RhoA inhibition, C3 exoenzyme protein (CT03; Cytoskeleton, Inc.) was resuspended and dialyzed into PBS and injected at 1 mg/mL. Fifteen embryos were imaged and quantifications in Fig. 4 D are from 4 representative embryos. PBS was injected as a control. To depolymerize F-actin we resuspended CtyoD (Enzo Life Sciences) at either 5 mg/mL or .125 mg/mL in DMSO or LatB (Enzo Life Sciences) in DMSO (5 mg/mL). Colchicine and taxol were both resuspended at 5 mg/mL in DMSO. Embryos were imaged 3-5 minutes after injection. For live injection of colchicine, embryos were mounted ventral side down onto a No.1 Coverslip. The embryo was pierced with the needle on the confocal microscope and injected after tissue folding had initiated during live imaging. To PFA fix embryos after injection, embryos were mounted onto a coverslip (No. 1.5) coated with a strip of glue. After injection, coverslips were placed in to a petri dish filled with heptane for ~45 seconds to remove embryos from the glue. Embryos in heptane were transferred to a 10% paraformaldehyde solution in 0.1 M phosphate buffer at pH 7.4 with 50% heptane and fixed for 30 minutes and manually devitellinized.

### Online Supplemental Material

Fig. S1 shows the location of centrosomes, verification of *patronin* RNAi, and the effect on microtubule organization after both genetic and pharmacological disruptions. Fig. S2 shows the effect of F-actin depolymerization on Patronin localization. Fig. S3 shows the phenotype of *patronin*-RNAi and the effect of taxol on microtubule organization. Fig. S4 shows that microtubule disruption does not affect the ability of actomyosin to reattach and does not affect medioapical ROCK polarity. In addition, the results of the live colchicine injection and the effect of microtubule disruption on E-cadherin polarity at late stages are shown. Video 1 shows the dynamic changes in localization of Patronin in the mesoderm and ectoderm. Video 2 shows the co-localization of myosin with dense patches of CLIP170. Video 3 shows the coalescence of Patronin correlating with myosin pulses and their co-localization. Video 4 shows the phenotypes of microtubule drug injections on tissue invagination. Video 5 shows the dynamic myosin network separations after colchicine injection. Video 6 shows the effect on the F-actin meshwork after taxol injection. Table S1 lists genotypes for all fly stocks and crosses used in this study.

## Acknowledgments

We would like to thank members of the Martin lab for their helpful comments and suggestions on the project and the manuscript. We thank Frank Mason and Iain Cheeseman for their helpful comments on a draft of this manuscript. We would also like to thank Mark Biggin (Berkeley Lab), Mark Peifer, Jordan Raff (University of Oxford), Ron Vale (UCSF), Steve Wasserman (UCSD), the Bloomington Stock Center, and TRiP at Harvard Medical School (National Institutes of Health/National Institute of General Medical Sciences R01-GM084947) for fly stocks and antibodies used in this study. This work was supported by grant R01GM105984 to A. C. Martin from the National Institute of General Medical Sciences. The authors declare no competing financial interests.

## Author contributions

C. S. Ko and A. C. Martin conceptualized the project and designed experiments. C. S. Ko and A. C. Martin performed the experiments. C. S. Ko and V. Tserunyan analyzed the data. C. S. Ko and A. C. Martin wrote the manuscript. All authors reviewed and approved the final version of the manuscript.

---

**Video 1. Patronin::GFP forms a medioapical focus in mesoderm cells.** Embryos expressing Patronin::GFP (green) and Gap43::mCH (magenta) shown with the midline of the mesoderm centered (top) or slightly turned (bottom) to show the ectoderm. Images were acquired every 44 seconds (top) or 40 seconds (bottom) and videos are displayed at 10 frames per second. Bars, 15 μm.

**Video 2. GFP::CLIP170 puncta co-localize with apical myosin.** Embryo expressing GFP::CLIP170 (green) and Myo::mCH (magenta). Images were acquired every 1.9 seconds and video is displayed at 15 frames per second. Bars, 15 μm.

**Video 3. Patronin medioapical foci in mesoderm cells form by myosin contraction.** Embryo expressing Patronin::GFP (green) and Myo::mCH (magenta). Images were acquired every 1.9 seconds and video is displayed at 15 frames per second. Bars, 15 μm.

**Video 4. Colchicine and taxol disrupt mesoderm invagination but does not interfere with apical constriction initiation.** Embryos expressing Myo::GFP (green) and Gap43::mCH (magenta) were injected with DMSO (top), colchicine (5 mg/mL; middle), or taxol (5 mg/mL; bottom). Images were acquired every 6.4 seconds and videos are displayed at 15 frames per second. Bars, 15 μm.

**Video 5. Microtubule disruption destabilizes intercellular actomyosin attachments.** Embryo expressing Myo::GFP was injected with colchicine (5 mg/mL). Images were acquired every 24 seconds and video is displayed at 15 frames per second. Bar, 15 μm.

**Video 6. The F-actin cortex exhibits longer-lived fractures and separations from junctions after microtubule disruption.** Embryo expressing Utr::GFP was injected with taxol (5 mg/mL). Images were acquired every 19.6 seconds and video is displayed at 15 frames per second. Bar, 15 μm.

## Supplemental Material

**Table S1. List of genotypes of fly stocks used in this study as well as specific crosses that generated the data**

**Figure S1.**
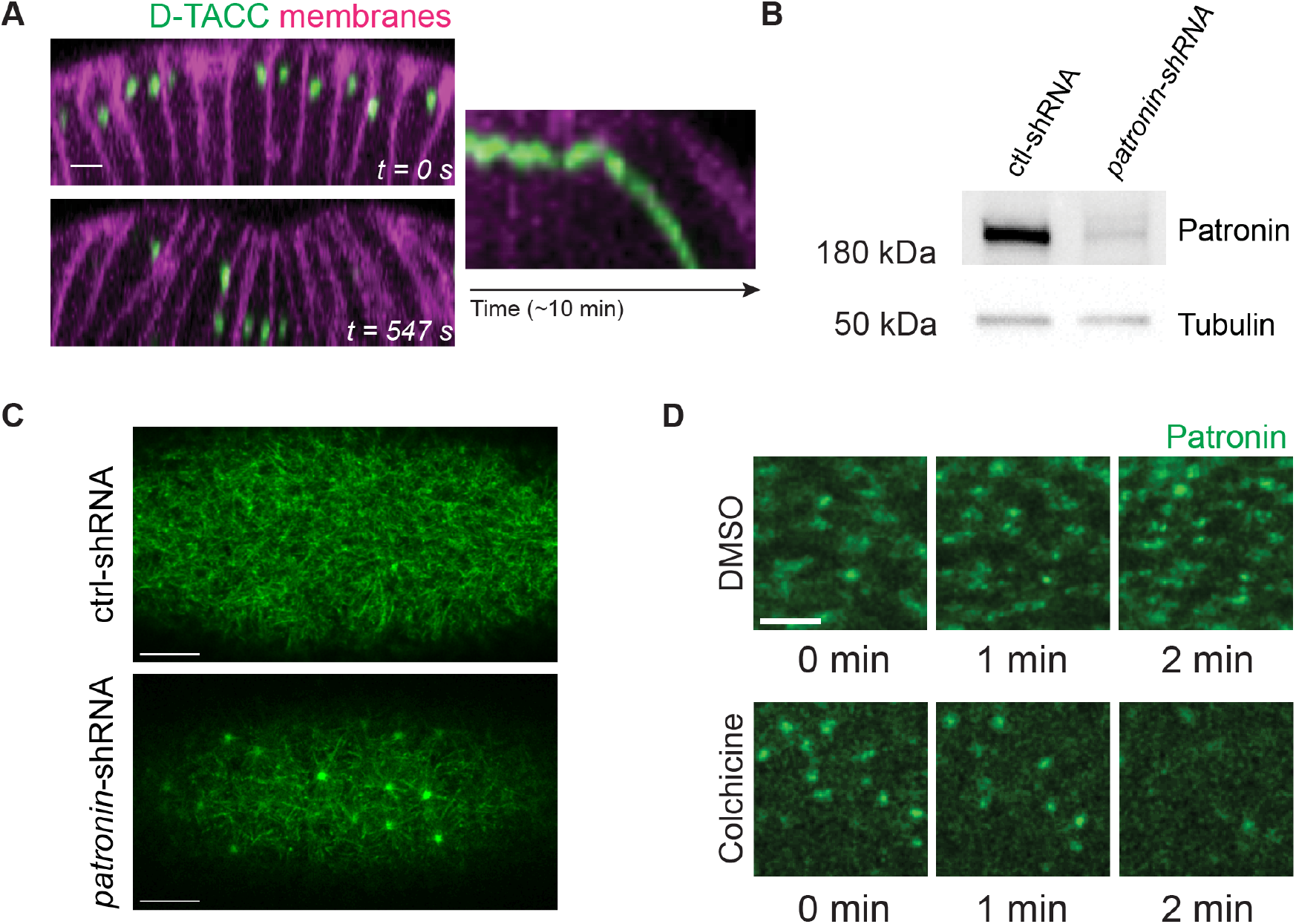
Medioapical Patronin foci are not centrosomes. (A) Time-lapse images showing an apical-basal cross-section from a live embryo expressing DTACC::GFP (green, a marker for centrosomes; Gergely et al., 2000) and Gap43::mCH (magenta). A kymograph using a line drawn along the apical-basal axis of a cell at the midline is shown on the right. (B) Lysates from control *rhodopsin-3*-shRNA and *patronin*-shRNA flies probed with Patronin antibody serum (gift from R. Vale). Anti-α-Tubulin was used as a loading control. (C) Microtubule organization is disrupted after Patronin depletion. Images are single apical slices from a live movie of a representative Rhodopsin-3 control RNAi (top) *patronin*-RNAi (bottom) embryo expressing GFP::CLIP170 (D). Patronin::GFP localization depends on an intact microtubule cytoskeleton. Images are a montage from a live movie of an embryo expressing Patronin::GFP and injected with either DMSO (top) or Colchicine (5 mg/mL; bottom). Scale bars, 15 μm (**C**), 5 μm (**A and D**).

**Figure S2.**
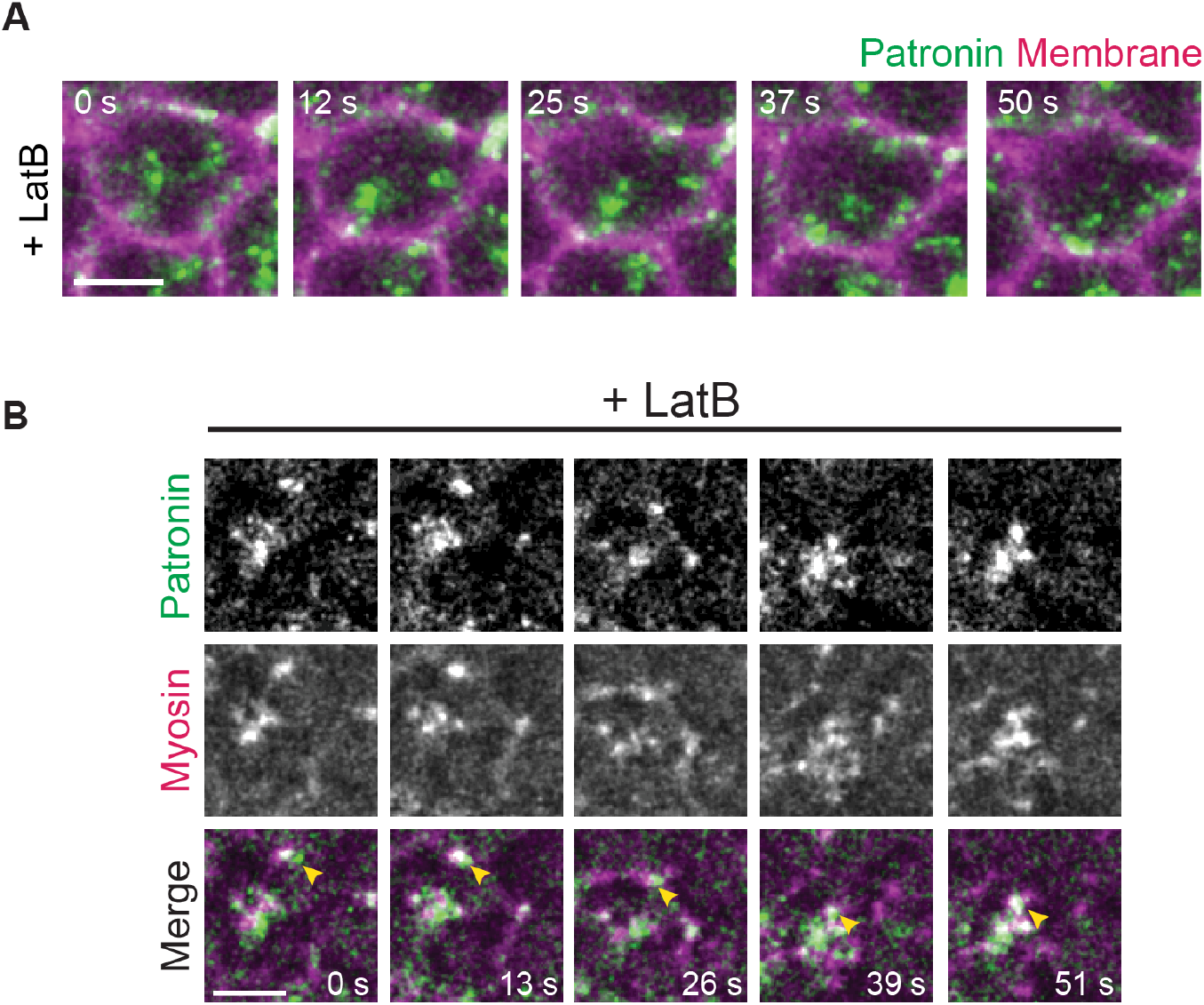
CytoD and LatB affect the organization of apical microtubules. (A) Patronin foci fragment into puncta after LatB injection. Time-lapse images are maximum intensity Z-projections of a representative embryo expressing Patronin::GFP and Gap43:mCH injected with Latrunculin B (5 mg/mL in DMSO). (B) Patronin puncta co-localize with myosin puncta after F-actin network fragmentation with LatB (yellow arrowheads). Time-lapse images are maximum intensity Z-projections of a representative embryo expressing and Patronin::GFP and Myo::mCH injected with LatB (5 mg/mL in DMSO). Scale bars, 5 μm.

**Figure S3.**
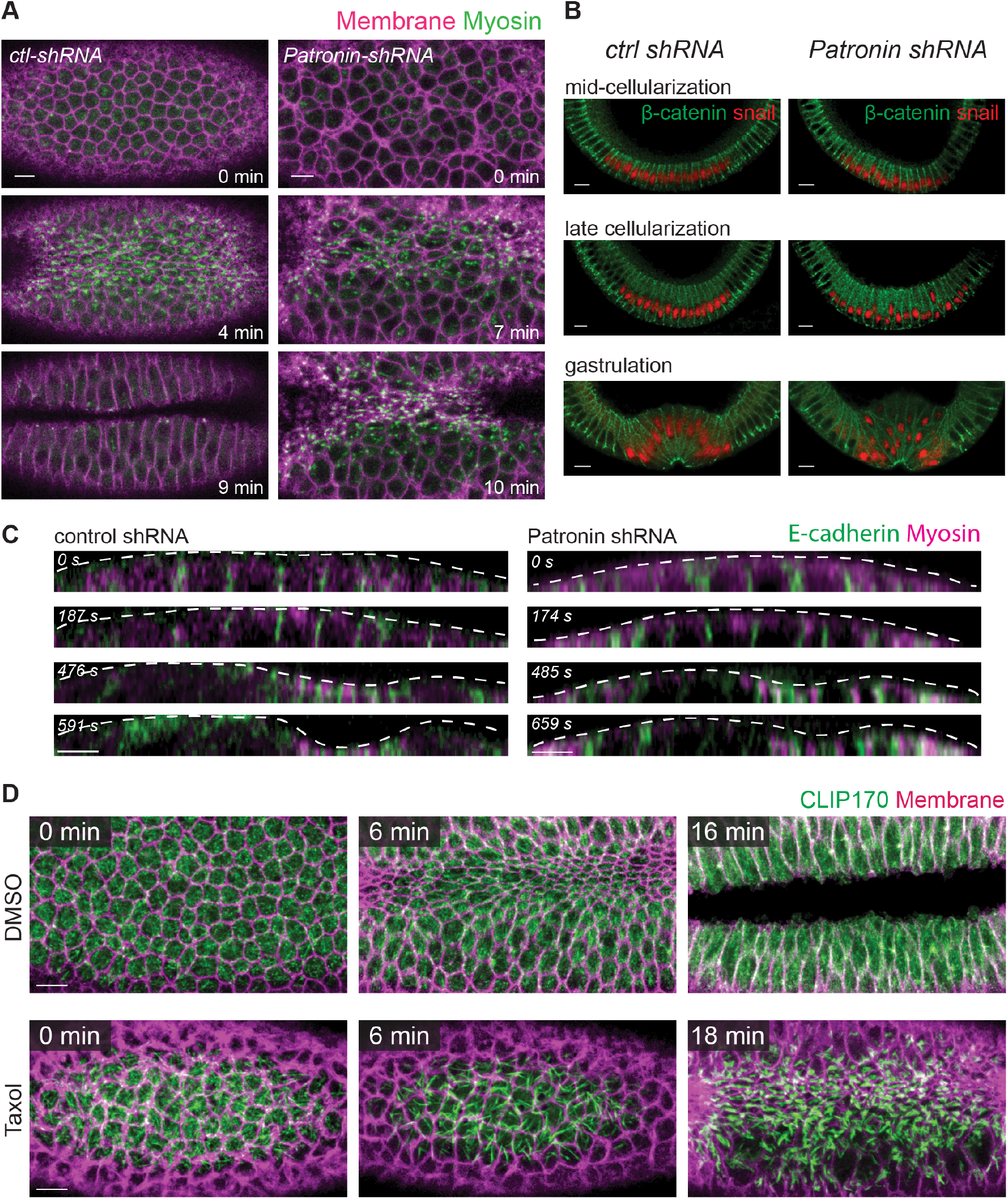
Patronin depletion disrupts folding, but not apical adherens junctions. (A) Depleting Patronin disrupts folding, despite apical myosin activation and apical constriction initiation. Time-lapse images are maximum intensity Z-projections from live embryos expressing control-shRNA (left) or *patronin*-shRNA (right) and Myo::GFP (apical surface) and Gap43::mCH (sub-apical section). The phenotype was observed in 5 out of 12 embryos imaged from this cross. (B) In mesoderm cells, adherens junctions exhibit apical shift after Patronin depletion. Images are apical-basal cross-sections from embryos fixed at different developmental stages stained for β-catenin (Armadillo) and Snail. (C) Apical adherens junctions are unaffected by Patronin depletion. Time-lapse images show apical-basal cross sections from representative live embryos expressing *control*-shRNA (left) or *patronin*-shRNA (right) and E-cadherin::GFP and Myo::mCH. Thirty-two embryos were imaged in total. (D) Taxol injection causes dense microtubule bundles across the apical surface. Time-lapse images are maximum intensity Z-projections from live embryos expressing GFP::CLIP170 (apical surface) and Gap43::mCH (sub-apical section) injected with either DMSO (top) or taxol (5 mg/mL; bottom). Scale bars, 10 μm (**A, C, D**), 5 μm (**B**).

**Figure S4.**
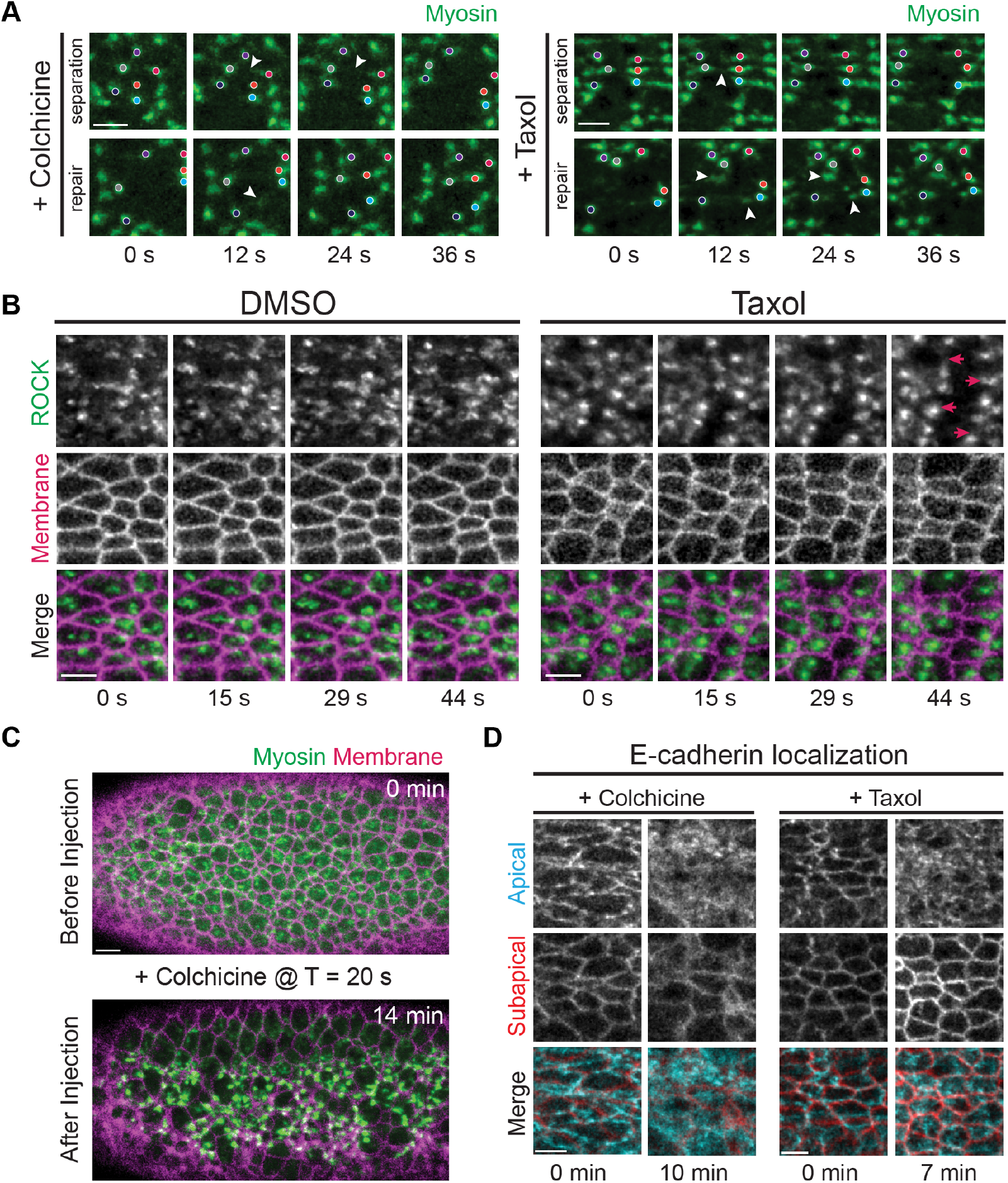
Disrupting microtubules causes actomyosin network separations from adherens junctions. (A) Myosin separation events are dynamic and attachments are reestablished following separation. Time-lapse images are maximum intensity Z-projections of apical Myo::GFP in embryos injected with colchicine or taxol. Individual myosin patches are labeled with colored dots. Fiber-like structures between myosin patches that either break or re-form during repair are shown by white arrowheads. (B) Taxol does not affect ROCK polarity, but causes separation between ROCK foci and junctions. Time-lapse images are maximum intensity Z-projections from live embryos expressing GFP::ROCK and Gap43::mCH injected with DMSO or taxol (5 mg/mL). Red arrows indicate the direction in which medioapical ROCK foci separate from cell junctions. (C) Microtubules were acutely inhibited by injecting embryo with colchicine ~20 seconds after start of imaging. Images are maximum intensity Z-projections from a live embryo expressing Myo::GFP and Gap43::mCH. (D) E-cadherin eventually loses junctional polarity over time. Time-lapse images showing a single apical (cyan) and sub-apical (red) slice of representative embryos expressing E-cadherin::GFP injected with colchicine (5 mg/mL) (left) and taxol (5 mg/mL) (right). Scale bars, 10 μm (**C**), 5 μm (**A-B, D**).

